# Genes with high network connectivity are enriched for disease heritability

**DOI:** 10.1101/442582

**Authors:** Samuel S. Kim, Chengzhen Dai, Farhad Hormozdiari, Bryce van de Geijn, Steven Gazal, Yongjin Park, Luke O’Connor, Tiffany Amariuta, Po-Ru Loh, Hilary Finucane, Soumya Raychaudhuri, Alkes L. Price

**Affiliations:** Department of Electrical Engineering and Computer Science, Massachusetts Institute of Technology, Cambridge, MA; Department of Epidemiology, Harvard T.H. Chan School of Public Health, Boston, MA; Program in Medical and Population Genetics, Broad Institute of MIT and Harvard, Cambridge, MA; Department of Computer Science, Harvard University, Cambridge, MA; Department of Biostatistics, Harvard T.H. Chan School of Public Health, Boston, MA; Program in Bioinformatics and Integrative Genomics, Harvard University, Cambridge, MA; Division of Genetics, Department of Medicine, Brigham and Women’s Hospital and Harvard Medical School, Boston, MA

## Abstract

Recent studies have highlighted the role of gene networks in disease biology. To formally assess this, we constructed a broad set of pathway, network, and pathway+network annotations and applied stratified LD score regression to 42 independent diseases and complex traits (average *N*=323K) to identify enriched annotations. First, we constructed annotations from 18,119 biological pathways, including 100kb windows around each gene. We identified 156 pathway-trait pairs whose disease enrichment was statistically significant (FDR < 5%) after conditioning on all genes and on annotations from the baseline-LD model, a stringent step that greatly reduced the number of pathways detected; most of the significant pathway-trait pairs were previously unreported. Next, for each of four published gene networks, we constructed probabilistic annotations based on network connectivity using closeness centrality, a measure of how close a gene is to other genes in the network. For each gene network, the network connectivity annotation was strongly significantly enriched. Surprisingly, the enrichments were fully explained by excess overlap between network annotations and regulatory annotations from the baseline-LD model, validating the informativeness of the baseline-LD model and emphasizing the importance of accounting for regulatory annotations in gene network analyses. Finally, for each of the 156 enriched pathway-trait pairs, for each of the four gene networks, we constructed pathway+network annotations by annotating genes with high network connectivity to the input pathway. For each gene network, these pathway+network annotations were strongly significantly enriched for the corresponding traits. Once again, the enrichments were largely explained by the baseline-LD model. In conclusion, gene network connectivity is highly informative for disease architectures, but the information in gene networks may be subsumed by regulatory annotations, such that accounting for known annotations is critical to robust inference of biological mechanisms.

## Introduction

Human diseases and complex traits are heritable and highly polygenic, potentially involving a large number of disease genes connected by dense cellular networks (Chen et al., 2008; Barabási et al., 2011; Menche et al., 2015; Visscher et al., 2017; Boyle et al., 2017). Recent work has employed several approaches to infer gene interaction networks, including protein-protein interaction networks (Szklarczyk et al., 2014; Chatr-Aryamontri et al., 2017; Li et al., 2017), tissue-specific co-expression networks (Greene et al., 2015; Saha et al., 2017), and tissue-specific regulatory networks (Marbach et al., 2016; Sonawane et al., 2017). An appealing extension of traditional genome-wide association studies (GWAS) is to identify genes and gene pathways associated with disease by leveraging gene networks and network connectivity between disease genes (Köhler et al., 2008; Vanunu et al., 2010; Lee et al., 2011; Califano et al., 2012; Taşan et al., 2015; Greene et al., 2015; Marbach et al., 2016; Peters et al., 2017; Cowen et al., 2017; Yoon et al., 2018). However, despite considerable progress on inferring gene interaction networks and applying newly developed methods for network connectivity-informed GWAS to identify specific genes and gene pathways associated to disease, an overall assessment and interpretation of the contribution of gene networks to the genetic architecture of disease has remained elusive. In particular, the extent to which this contribution can be explained by disease enrichments of known functional annotations (Consortium et al., 2012; Maurano et al., 2012; Trynka et al., 2013; Kundaje et al., 2015; Finucane et al., 2015; Gazal et al., 2017) is unknown.

Here, we sought to answer three questions. First, what is the contribution of disease-associated gene pathways (Segrè et al., 2010; Pers et al., 2015; de Leeuw et al., 2015, 2016; Boyle et al., 2017; Pardiñas et al., 2018; Yoon et al., 2018; Wray et al., 2018; Savage et al., 2018; Zhu and Stephens, 2018; Nagel et al., 2018) to disease heritability, irrespective of network connectivity. Second, what is the contribution of genes with high network connectivity in known gene networks (Greene et al., 2015; Saha et al., 2017; Li et al., 2017; Sonawane et al., 2017) to disease heritability. Third, what is the contribution of genes with high network connectivity to disease-associated gene pathways to disease heritability. We hypothesized that genes with high network connectivity to disease-associated gene pathways would be more informative than gene pathways or gene networks alone, as information in gene networks is orthogonal to information in gene pathways.

To answer these questions, we constructed a broad set of pathway, network, and pathway+network annotations. The pathway annotations were constructed from known gene pathways by including 100kb windows around each gene; the network annotations were constructed by quantifying network connectivity using closeness centrality, a measure of how close a gene is to other genes in the network (Boyle et al., 2018; Li et al., 2018); and the pathway+network annotations were constructed by annotating genes with high network connectivity to the input pathway, again quantified using closeness centrality. We applied stratified LD score regression (Finucane et al., 2015; Gazal et al., 2017) to quantify the contribution of the pathway, network, and pathway+network annotations to disease heritability. We conditioned our analyses on all genes and on the baseline-LD model, which includes a broad set of coding, conserved, regulatory and LD-related annotations (Gazal et al., 2017). In each case, we compared results before and after conditioning on the baseline-LD model, to assess the extent to which the disease enrichments that we identified could be explained by known functional annotations.

## Results

### Overview of methods

We provide an overview of the methods and data used in our three main analyses, in which we assessed the enrichment of disease/trait heritability in pathway, network, and pathway+network based genomic annotations, respectively. First, in our pathway analyses, we considered 18,119 sets of protein-coding genes from five sources: 2,118 biological pathways from the BioSystem (BS) database (Geer et al., 2010), 1,927 biological pathways from the Pathway Commons (PC) database (Cerami et al., 2011), 7,209 protein-protein interaction gene sets from the InWeb database (Li et al., 2017), 3,903 mouse phenotype gene sets from the Mouse Genome Informatics (MGI) database (Eppig et al., 2014) (i.e. sets of genes whose orthologs are associated to mouse phenotypes), and 2,961 gene ontology gene sets from the Genome Ontology (GO) database (Ashburner et al., 2000) (see Methods). We refer to these 18,119 gene sets as “pathways.” We restricted to pathways with 10-500 genes, as in previous pathway enrichment studies (Pers et al., 2015; Zhu and Stephens, 2018). The complete list of pathways is provided in Table S1, and a histogram of the number of genes in pathways from each of the five sources is provided in Figure S1. We applied stratified LD score regression (S-LDSC) (Finucane et al., 2015; Gazal et al., 2017) to binary pathway annotations constructed using ±100kb windows around each gene, as in previous work (Finucane et al., 2018; Zhu and Stephens, 2018). We analyzed publicly available GWAS summary statistics from 42 independent diseases and complex traits, including 30 UK Biobank traits (see URLs and Table S2; average *N* = 323K) to evaluate the contribution of each pathway annotation to disease/trait heritability. We conditioned on the 75 functional annotations from the baseline-LD model (Gazal et al., 2017) (which includes a broad set of coding, conserved, regulatory and LD-related annotations), as well as an all-genes annotation representing the set of all 23,987 protein-coding genes (±100kb). For each of 760,869 pathway-trait pairs (roughly 18,119 pathways x 42 traits; see Methods), we assessed the statistical significance of the pathway annotations standardized effect size *τ*\* (defined as the proportionate change in per-SNP heritability associated to a one standard deviation increase in the value of the annotation, conditioned on other annotations included in the model (Gazal et al., 2017)) based on global false discovery rate (FDR) < 5%; we also computed the enrichment, defined as the proportion of heritability divided by the proportion of SNPs. Unlike enrichment, *τ*\* quantifies effects that are unique to the focal annotation (see Methods).

Second, in our network analyses, we considered four gene networks: two co-expression networks (Saha (Saha et al., 2017) and Greene (Greene et al., 2015)), one protein-protein interaction network (InWeb (Li et al., 2017)), and one regulatory network (Sonawane (Sonawane et al., 2017)). A gene network is defined by an edge weight (which we normalized to lie between 0 and 1) for each pair of genes, representing their connectivity in the network; we note that the Sonawane regulatory network contains nonzero edge weights only for pairs of genes in which at least one of the genes is a known transcription factor. We further note that the Greene, Saha and Sonawane networks include tissue-specific networks for 144, 18 and 38 tissues, respectively; the choice of which tissue-specific network to use for each disease/trait is described below (see Enrichment of disease heritability in network annotations). For each network analyzed, we computed the closeness centrality (Özgür et al., 2008) for each protein-coding gene (a measure of network connectivity, i.e. how connected the gene is to other protein-coding genes in the network) and constructed a probabilistic annotation by linearly transforming connectivity values to lie between 0 to 1 and including ±100kb windows as above; we also considered other connectivity metrics (see Methods). For each of 168 network-trait pairs (4 networks x 42 traits), we applied S-LDSC and assessed the statistical significance of the network annotations *τ*\* conditioned on other annotations; we also computed the enrichment, which naturally extends to probabilistic annotations (Hormozdiari et al., 2018).

Third, in our pathway+network analyses, we considered 156 significant pathway-trait pairs (from our pathway analyses) and four gene networks. For each input pathway and each input network, we defined a set of candidate genes based on the genes in the input pathway and their one-degree neighbors in the input network; defined a subnetwork by restricting the input network to this set of candidate genes; computed the closeness centrality (Özgür et al., 2008) for each candidate gene in that subnetwork; and constructed a probabilistic annotation by linearly transforming connectivity values to lie between 0 to 1 and including ±100kb windows as above. For each of 590 (pathway-trait,network) pairs (122 pathway-trait pairs for Saha + 156 pathway-trait pairs x 3 other networks; see Methods), we applied S-LDSC and assessed the statistical significance of the pathway+network annotations *τ*\* conditioned on other annotations; we also computed the enrichment.

We have publicly released all pathway, network and pathway+network annotations, LD scores, and S-LDSC results (see URLs).

### Enrichment of disease heritability in pathway annotations

We sought to identify pathways that are enriched for disease heritability. We applied S-LDSC to 760,869 pathway-trait pairs, spanning 18,119 pathways from five sources (Table S1 and Figure S1) and 42 independent diseases and complex traits, including 30 UK Biobank traits (average *N* = 323K; Table S2). We identified 156 pathway-trait pairs (spanning 141 pathways and 34 traits) that were significantly enriched after conditioning on the baseline-LD model and the all-genes annotation (FDR<5% for positive *τ*\*). Complete results for three representative traits (Crohns disease, rheumatoid arthritis, and schizophrenia (Boyle et al., 2017)) are reported in Figure 1, and complete results for all traits are reported in Table S3. The top pathway (most significant *τ*\*) for each of the 34 traits is reported in Table 1, and the complete set of 156 significant pathway-trait pairs is reported in Table S4. Genes in the 141 enriched pathways had a larger gene size (92kb on average) compared to all protein-coding genes (58kb) and genes in all pathways (76kb) (see Table S5). We meta-analyzed the 156 pathway-trait pairs using random-effect meta-analysis (analogous to previous work (Finucane et al., 2015; Gazal et al., 2017; Hormozdiari et al., 2018)). Both the enrichment (4.13, s.e. = 0.12; P = 4.74e-158) and *τ*\* (0.15, s.e. = 0.0061; P=3.41e-131) were large and highly statistically significant (Table S6); we caution that these p-values are slightly inflated because we meta-analyzed across significant pathway-trait pairs only.

**Figure 1.**
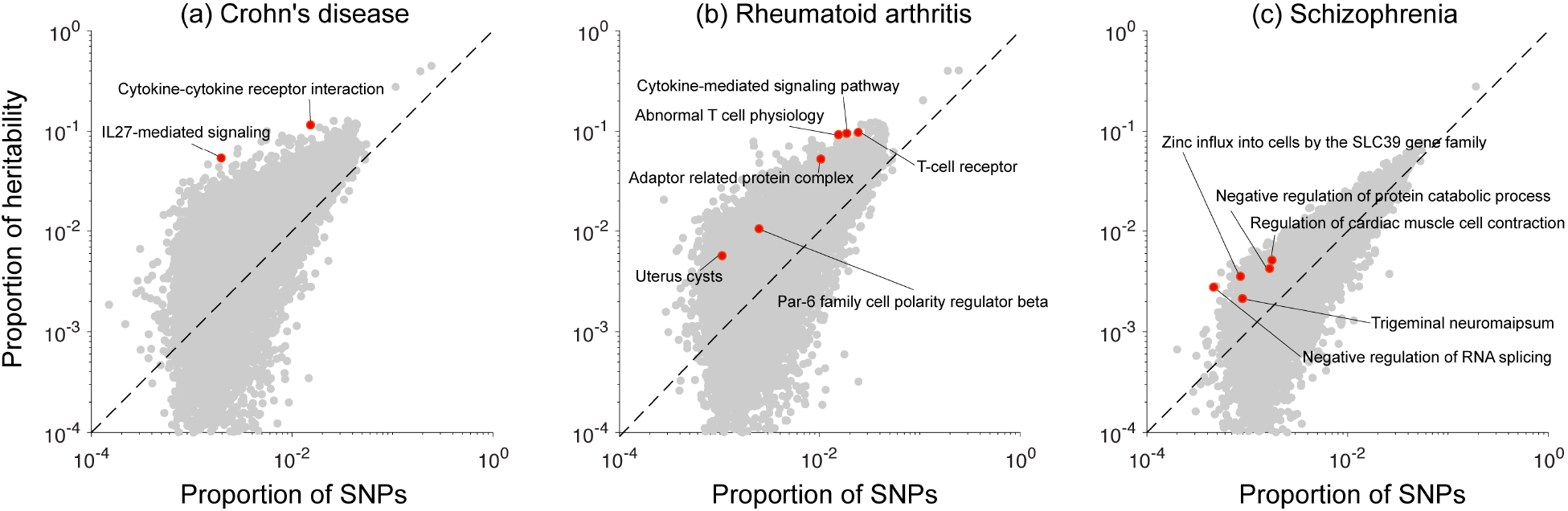
Enriched pathways for three representative traits. For (a) Crohns disease, (b) rheumatoid arthritis and (c) schizophrenia, we report the proportion of heritability explained and proportion of SNPs for each of 18,119 pathways analyzed. Red points indicate significantly enriched pathways (FDR < 5%), and grey points indicate non-significant pathways. Numerical results for all 42 diseases and complex traits are reported in Table S3.

**Table 1.** Top enriched pathway for each trait. We report the top enriched pathway (most significant *τ*\*) for each of 34 traits with at least one significantly enriched pathway. The first 23 traits (above the line) are UK Biobank traits. Enrichment of the melanin biosynthetic process pathway for skin color is consistent with previous studies (Low and Chen, 2011; Simeonov et al., 2013), and enrichment of the Absent corpus callosum pathway for years of education was reported in a previous genetic study (Okbay et al., 2016). The complete set of 156 significant pathway-trait pairs is reported in Table S4.

| Trait | Top enriched pathway | Database | #Genes | Enr. (s.e.) | $\tau^*$ (s.e.) |
| --- | --- | --- | --- | --- | --- |
| Age at menarche | SIX homeobox 6 | InWeb | 11 | 3.06 (0.45) | 0.06 (0.02) |
| Auto immune traits | Mismatch repair directed by MSH2:MSH6 | BS | 14 | 8.53 (2.05) | 0.28 (0.07) |
| BMI | Positive regulation of synapse maturation | GO | 11 | 3.34 (0.25) | 0.07 (0.01) |
| Dermatologic diseases | Aldo-keto reductase family 1 member E2 | InWeb | 12 | 7.51 (1.80) | 0.19 (0.05) |
| Eczema | Th17 cell differentiation | BS | 102 | 9.01 (1.56) | 0.64 (0.15) |
| Eosinophil count | Jak-STAT signaling pathway | BS | 152 | 7.67 (1.11) | 0.46 (0.11) |
| Forced vital capacity | Absent acrosome | MGI | 11 | 2.21 (0.31) | 0.04 (0.01) |
| Heel T Score | Skeletal system development | GO | 142 | 4.43 (0.60) | 0.31 (0.07) |
| Height | Abnormal trabecular bone morphology | MGI | 172 | 2.93 (0.39) | 0.20 (0.05) |
| Hypothyroidism | Cytokine-mediated signaling pathway | GO | 235 | 4.88 (0.78) | 0.42 (0.11) |
| Morning person | N-acetylglucosamine metabolic process | GO | 13 | 2.07 (0.26) | 0.03 (0.01) |
| Neuroticism | HSF1 activation | BS | 11 | 2.54 (0.27) | 0.04 (0.01) |
| Platelet count | Platelet activation | GO | 208 | 4.32 (0.62) | 0.30 (0.09) |
| Red blood cell count | Exogenous drug catabolic process | GO | 10 | 3.62 (0.63) | 0.08 (0.02) |
| Red blood cell distribution width | Decreased erythrocyte cell number | MGI | 206 | 4.61 (0.74) | 0.39 (0.11) |
| Respiratory/ear-nose-throat diseases | Abnormal T cell activation | MGI | 108 | 4.28 (0.75) | 0.26 (0.07) |
| Skin color | Melanin biosynthetic process | GO | 11 | 192.14 (55.37) | 5.90 (1.73) |
| Smoking status | Fibromodulin | InWeb | 17 | 2.53 (0.48) | 0.06 (0.02) |
| Systolic blood pressure | cGMP-PKG signaling pathway | BS | 164 | 3.41 (0.49) | 0.25 (0.07) |
| Type 2 diabetes | Glucuronidation | BS | 30 | 2.74 (0.32) | 0.06 (0.01) |
| Waist-hip ratio | Negative regulation of transcription | GO | 496 | 2.60 (0.22) | 0.19 (0.05) |
| White blood cell count | ERK cascade | PC | 15 | 4.65 (0.69) | 0.11 (0.03) |
| Years of education | Absent corpus callosum | MGI | 45 | 2.50 (0.33) | 0.08 (0.02) |
| Age first birth | Receptor signaling protein tyrosine kinase | GO | 10 | 3.62 (0.58) | 0.10 (0.02) |
| Anorexia | Activation of the AP-1 family of TF | BS | 10 | 5.64 (1.04) | 0.15 (0.03) |
| Autism spectrum | GABA-A receptor activity | GO | 15 | 4.01 (0.73) | 0.12 (0.02) |
| Coronary artery disease | Sodium-independent organic anion transport | GO | 12 | 7.05 (1.49) | 0.22 (0.04) |
| Crohn's disease | Cytokine-cytokine receptor interaction | BS | 266 | 7.59 (1.50) | 0.65 (0.18) |
| Depressive symptoms | Glycoprotein metabolic process | GO | 12 | 5.22 (1.10) | 0.15 (0.04) |
| LDL | Increased erythroblast number | MGI | 21 | 5.63 (1.15) | 0.18 (0.05) |
| Number children even born | Lipoxygenase pathway | GO | 12 | 9.81 (1.59) | 0.27 (0.04) |
| Rheumatoid arthritis | Par-6 family cell polarity regulator beta | InWeb | 16 | 4.31 (1.01) | 0.17 (0.04) |
| Schizophrenia | Trigeminal neuroma | MGI | 10 | 2.39 (0.31) | 0.03 (0.01) |
| Ulcerative colitis | FGFR2b ligand binding and activation | BS | 10 | 8.00 (1.97) | 0.18 (0.05) |

Our results include 8 pathway-trait pairs reported in previous genetic studies (see Table 1 and Table S4). These include “immune response” for both Crohn’s disease and ulcerative colitis (Jostins et al., 2012); “T-cell receptor”, “cytokine-mediated signaling pathway”, and “abnormal T-cell physiology” for rheumatoid arthritis (Okada et al., 2014); and “Absent corpus callosum” for years of education (Okbay et al., 2016). In addition, “melanin biosynthetic process” was overwhelmingly enriched for skin color (Table 1), consistent with the fact that genetic variants in constituent genes are strongly associated with skin pigmentation (e.g. *MC1R*) and other pigmentation traits (e.g. *TYR*, *OCA2*, *SLC45A2*) (Low and Chen, 2011; Simeonov et al., 2013).

Surprisingly, most pathway-trait pairs reported in recent studies (Boyle et al., 2017; Pardiñas et al., 2018; Wray et al., 2018; Savage et al., 2018; Zhu and Stephens, 2018; Nagel et al., 2018) using genome-wide polygenic methods (de Leeuw et al., 2015; Finucane et al., 2015; Zhu and Stephens, 2018) were not significant in our analysis. Specifically, we considered 95 significant pathway-trait pairs for the 6 traits (schizophrenia, Crohn’s disease, rheumatoid arthritis, neuroticism, intelligence, depressive symptoms), analyzed in five previous studies (Boyle et al., 2017; Pardiñas et al., 2018; Wray et al., 2018; Savage et al., 2018; Nagel et al., 2018), restricting to at most the top 20 pathways per trait per study. Only 15/95 pathway-trait pairs were significant in our primary analysis, after conditioning on the baseline-LD model and all-genes annotation (Table S7A). However, 67/95 were significant when we repeated the S-LDSC analysis conditioning on just the all-genes annotation and not the baseline-LD model (Table S7B). We obtained similar results for a recent unpublished study (Zhu and Stephens, 2018) (Table S7A-B). Enriched pathways that were fully explained by the baseline-LD model are likely due to factors that do not play a direct role in trait biology (de Leeuw et al., 2018). Thus, it is critical to account for known functional annotations in pathway analyses to correct for such factors.

Our results also highlight pathway-trait pairs that have not previously been reported but are consistent with known biology. These include “GABA-A receptor activity” with autism (Table 1), consistent with the finding that the brains of subjects with autism have altered expression of GABA receptors (Fatemi et al., 2009; Cellot and Cherubini, 2014); “Oncostatin M (OSM)” for ulcerative colitis (Table S4), consistent with the finding that inflamed intestinal tissues from patients with inflammatory bowel diseases contained higher expression of OSM, and OSM-deficient mice displayed significantly attenuated colitis (West et al., 2017); and “zinc influx into cells by the SLC39 gene family” with schizophrenia (Table S4), consistent with the finding that *SLC39A12A* expression in dorsolateral prefrontal cortex is associated with schizophrenia (Scarr et al., 2016; Takagishi et al., 2017).

We also analyzed three additional gene sets (distinct from the 18,119 pathways) reflecting genes under strong selection: ExAC (Lek et al., 2016) (genes strongly depleted for protein-truncating variants), Cassa (Cassa et al., 2017) (genes with strong selection against protein-truncating variants), and Samocha (Samocha et al., 2014) (genes strongly depleted for missense mutations), which were previously shown to be enriched for heritability in a meta-analysis across traits (Hormozdiari et al., 2018). We identified 13 significantly enriched gene set-trait pairs (7 for ExAC, 2 for Cassa, and 4 for Samocha) spanning 9 traits, after conditioning on the baseline-LD model and the all-genes annotation (FDR < 5% for positive *τ*\*; Table S8). In a meta-analysis of these 13 gene set-trait pairs, both the enrichment (1.57, s.e. = 0.053; P = 7.52e-33) and *τ*\* (0.13, s.e. = 0.0090; P=3.30e-48) were highly statistically significant (Table S6).

### Enrichment of disease heritability in network annotations

We sought to assess the hypothesis that genes with high network connectivity are enriched for disease heritability. We constructed probabilistic annotations based on closeness centrality for each of the Saha, Greene, InWeb, and Sonawane networks (Table S9; see Methods). We determined that closeness centrality was independent of gene size (*r* = –0.015-0.019) and exon proportion (*r* = –0.18-0.008; Figure S2). For the three networks that include tissue-specific networks, we selected the Saha-skin(sun exposed lower leg), Greene-thyroid and Sonawane-testis networks for our primary analyses, as these tissue-specific networks maximized the correlation of the resulting network annotations with H3K27ac (Table S10; see below). We note that different tissue-specific networks from the same source were only weakly correlated (*r* = 0.027-0.076 for Saha, 0.17-0.21 for Greene, 0.062-0.24 for Sonawane; Table S10).

We computed the excess overlap between each of the four network annotations and representative annotations from the baseline-LD model and determined that each of the network annotations had substantial excess overlap with regulatory annotations (e.g. 1.23-1.36 with H3K27ac, 1.35-1.66 with H3K9ac; Figure 2a and Table S11), substantially stronger than the excess overlap between the all-genes annotation and regulatory annotations (e.g. 1.11 with H3K27ac, 1.14 with H3K9ac; Figure 2a); this implies that, for each network, genes with high connectivity to other genes in the network are enriched for the presence of nearby regulatory marks, perhaps because they are regulated by many other genes. Correlations between these annotations produced similar conclusions (Table S12; larger correlation for H3K27ac than H3K9ac), although we consider excess overlap to be a more robust metric because the size of the network annotations varies from 1%-34% of SNPs. We also report excess overlap and correlations between baseline-LD model annotations in Table S13.

**Figure 2.**
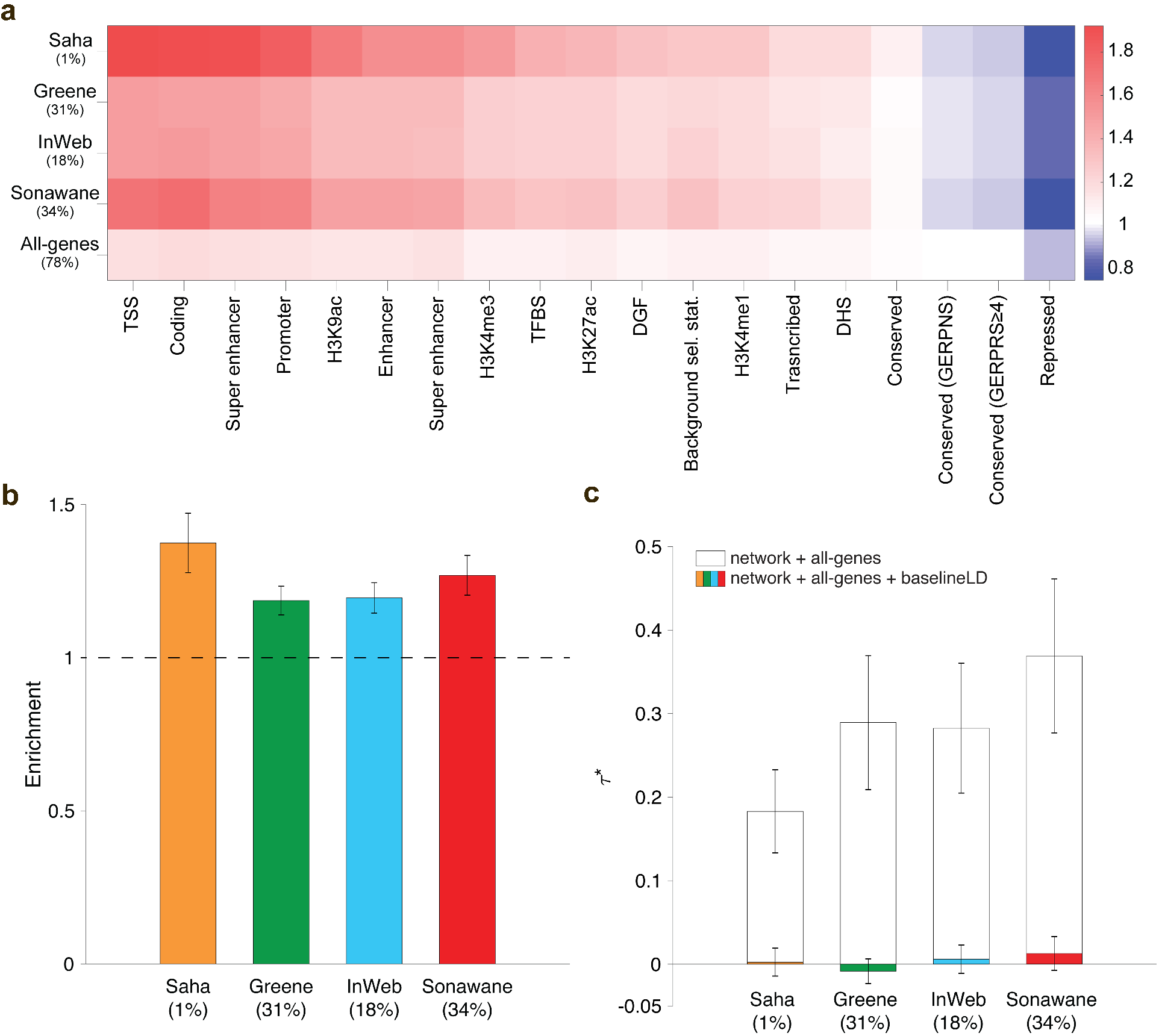
Heritability enrichment of network annotations. We report (a) excess (fold) overlap between network annotations and baseline-LD functional categories; (b) heritability enrichment of network annotations, meta-analyzed across 42 independent traits; and (c) *τ*\* values of network annotations, conditioned on either just the all-genes annotation, or the all-genes annotation and the baseline-LD model, meta-analyzed across 42 independent traits. In (b) and (c), the percentage under each bar indicates the proportion of SNPs in each annotation (defined for probabilistic annotations as the average value of the annotation), and error bars represent 95% confidence intervals. Numerical results for (a) are reported in Table S11, and numerical results for (b) and (c) are reported in Table S14.

For each of the four network annotations, we applied S-LDSC to the 42 independent diseases and complex traits, conditioning on the baseline-LD model and the all-genes annotation, and meta-analyzed the results across traits using random-effects meta-analysis. We identified strongly significant enrichments for each network annotation: 1.19 (s.e. = 0.024; P = 2.5e-30) to 1.37 (s.e. = 0.049; P = 1.6e-17) (Figure 2b and Table S14). However, estimates of *τ*\*, quantifying effects unique to the network annotations, were not significant (P = 0.21 to 0.77) (Figure 2c and Table S14). This implies that the enrichment signal in the network annotations (Figure 2b) is entirely explained by the excess overlap between the network and baseline-LD model annotations (Figure 2a); accordingly, when we repeated the S-LDSC analysis conditioning only on the all-genes annotation and not on the baseline-LD model, *τ*\* estimates were large and highly significant (Figure 2c). We repeated the S-LDSC analysis conditional on one annotation from the baseline-LD model at a time and confirmed that regulatory annotations (including histone marks and transcription factor binding sites) substantially reduced estimates of *τ*\* (Table S15). On the one hand, these findings represent a negative result for efforts to improve upon the baseline-LD model. On the other hand, these findings provide a strong validation of the baseline-LD model, in that the information about diseases/traits in annotations from other sources that broadly reflect the action of gene regulation are fully captured by the baseline-LD model.

We performed several secondary analyses. We repeated the S-LDSC analysis for the 3 tissue-specific networks (Saha, Greene, Sonawane) using the most biologically relevant tissue (as inferred from heritability enrichment of tissue-specific specifically expressed gene and chromatin annotations (Finucane et al., 2018)) for 10 blood-related traits (see Table S2 for list of traits and tissues); we obtained similar results, including significant enrichments for all three tissue-specific networks (Greene et al., 2015; Saha et al., 2017; Sonawane et al., 2017) but non-significant *τ*\* conditional on the baseline-LD model (Table S16). We further repeated the S-LDSC analysis for two tissue-specific networks (Greene et al., 2015; Sonawane et al., 2017) that include brain, for 8 brain-related traits; we obtained non-significant enrichments and non-significant *τ*\* conditional on the basline-LD model (Table S16). We also obtained similar results when we repeated the S-LDSC analysis using six connectivity metrics other than closeness centrality (see Methods; Table S17), using window sizes other than 100kb (10kb and 1Mb; Table S18), and constructing network annotations using diffusion state distance (DSD)-transformed networks (Cao et al., 2013) (Table S19).

### Enrichment of disease heritability in integrated pathway+network annotations

We sought to assess the hypothesis that genes with high network connectivity to genes in enriched pathways are also enriched for disease heritability; these enriched pathways serve as relevant, trait-specific signals that network analysis often lacks. For each of four networks and 141 enriched pathways, we constructed probabilistic pathway+network annotations based on closeness centrality within the subnetwork consisting of input genes and their one-degree neighbors. As before, for the three networks that include tissue-specific networks, we selected the Saha-skin(sun exposed lower leg), Greene-thyroid and Sonawane-testis networks for our primary analyses.

For each network, we computed the excess overlap between the 141 pathway+network annotations and representative annotations from the baseline-LD model and averaged results across the 141 pathways. We observed higher excess overlap with regulatory annotations (e.g. 1.27-1.35 with H3K27ac, 1.42-1.70 with H3K9ac; Figure 3a and Table S11) than the analogous excess overlaps for network annotations (Figure 2a); this implies that, for each network, genes with high network connectivity to genes in input pathways are enriched for the presence of nearby regulatory marks, perhaps because they are regulated by many other genes. Correlations between network, pathway+network and baseline-LD model annotations are reported in Table S12. We determined that pathway+network annotations were often correlated with network annotations, particularly for the corresponding network (from *r* = 0.12 for Saha to *r* = 0.93 for Greene). We further determined that pathway+network annotations constructed using the Greene, InWeb and Sonawane networks were moderately correlated (*r* = 0.27-0.42), whereas pathway+network annotations constructed using the Saha network were more distinct (*r* = 0.05-0.06), primarily due to its small annotation size (Table S12).

**Figure 3.**
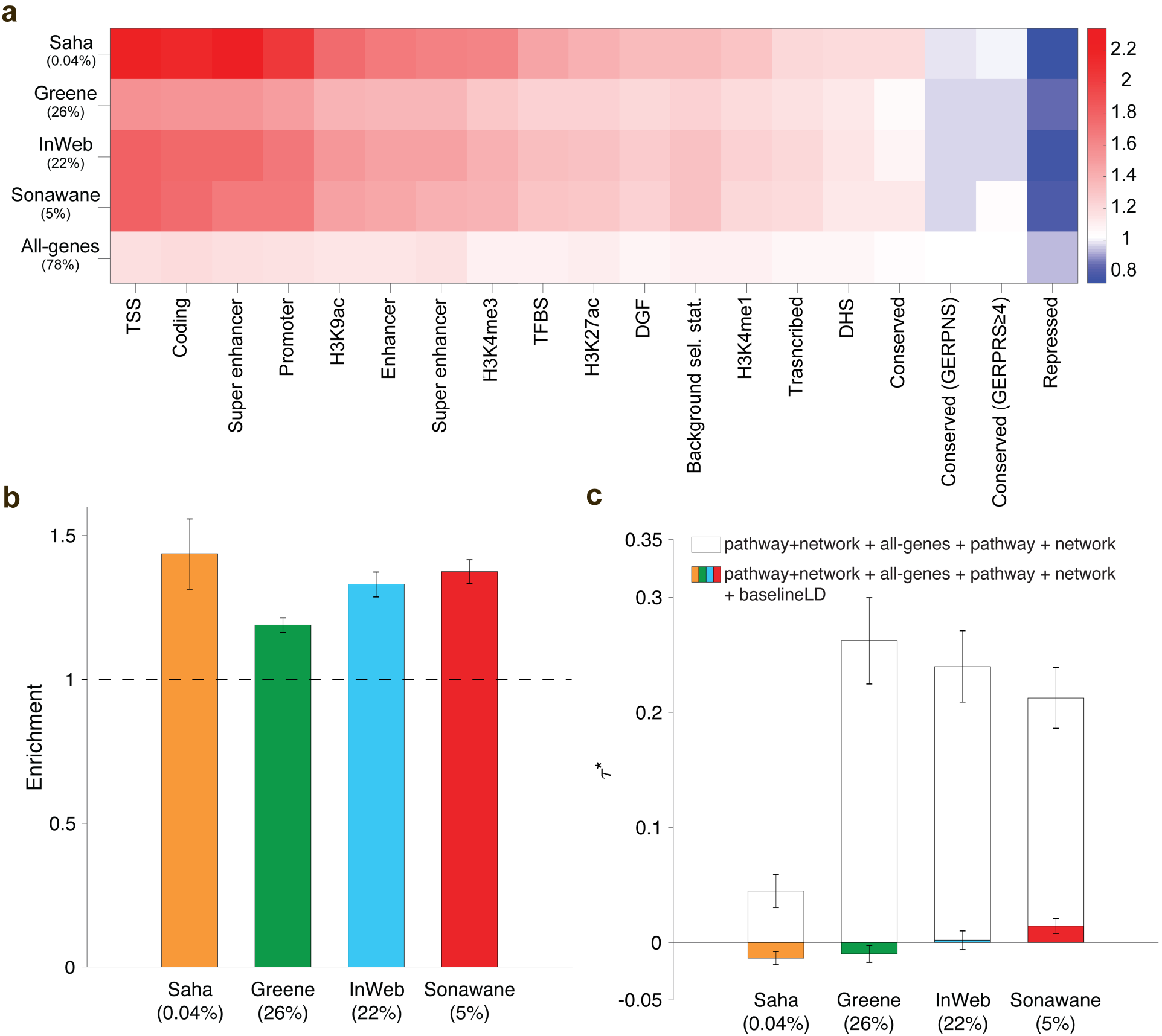
Heritability enrichment fof pathway+network annotations. We report (a) excess (fold) overlap between pathway+network annotations (averaged across up to 156 pathway-trait pairs); (b) heritability enrichment of pathway+network annotations, meta-analyzed across up to 156 pathway-trait pairs; and (c) *τ*\* values of pathway+network annotations, conditioned on either just the all-genes annotation and the corresponding pathway and network annotations, or the baseline-LD model as well, meta-analyzed across up to 156 pathway-trait pairs. In (b) and (c), the percentage under each bar indicates the proportion of SNPs in each annotation (defined for probabilistic annotations as the average value of the annotation), and error bars represent 95% confidence intervals. Numerical results for (a) are reported in Table S11, and numerical results for (b) and (c) are reported in Table S20.

For each of 590 (pathway-trait,network) pairs (122 pathway-trait pairs for Saha + 156 pathway-trait pairs × 3 other networks; see Methods), we applied S-LDSC to the resulting pathway+network annotation and the corresponding trait, conditioning on the baseline-LD model, the all-genes annotation, and the corresponding pathway and network annotations, and meta-analyzed the results for each network using random-effects meta-analysis. We identified strongly significant enrichments for all of our pathway+network annotations: 1.19 (s.e. = 0.01; P = 1.5e-49) to 1.44 (s.e. = 0.06; P = 3.8e-12) (Figure 3b and Table S20). However, estimates of *τ*\*, quantifying effects unique to the network annotations, were not significant or only weakly significant (P = 0.62 to 4.5e-6) (Figure 3c and Table S20). Once again, this implies that the enrichment signal in the pathway+network annotations is entirely explained by the excess overlap between the pathway+network and baseline-LD model annotations; accordingly, when we repeated the S-LDSC analysis conditioning only on the all-genes annotation and the corresponding pathway and network annotations, and not on the baseline-LD model, *τ*\* estimates were large and highly significant, except for the Saha network (Figure 3c). We repeated the S-LDSC analysis conditional on one annotation from the baseline-LD model at a time, and determined that inclusion of regulatory annotations (including histone marks and transcription factor binding sites) substantially reduced estimates of *τ*\* (Table S21).

We assessed whether genes in enriched pathways have higher network connectivity than other genes. For each of the four gene networks, for each of the 141 enriched pathways, we assessed both the network connectivity within the pathway and the network connectivity between the pathway and interacting genes (one-degree neighbors) outside the pathway, as compared to 10,000 null pathways with the same number of genes, each randomly sampled from a randomly chosen pathway from the full set of 18,119 pathways (see Methods). We assessed network connectivity using the sum of edge weights between genes. For each network, we averaged results across pathways. We determined that genes in enriched pathways have higher network connectivity within the pathway (1.43x−7.60x more edges; Figure 4 and Table S22), but do not necessarily have higher network connectivity with interacting genes outside the pathway (0.69x−1.56x; Table S22); we note that there is no significant difference in the number of interacting genes between the 141 enriched pathways and the 10,000 null pathways (Table S22).

**Figure 4.**
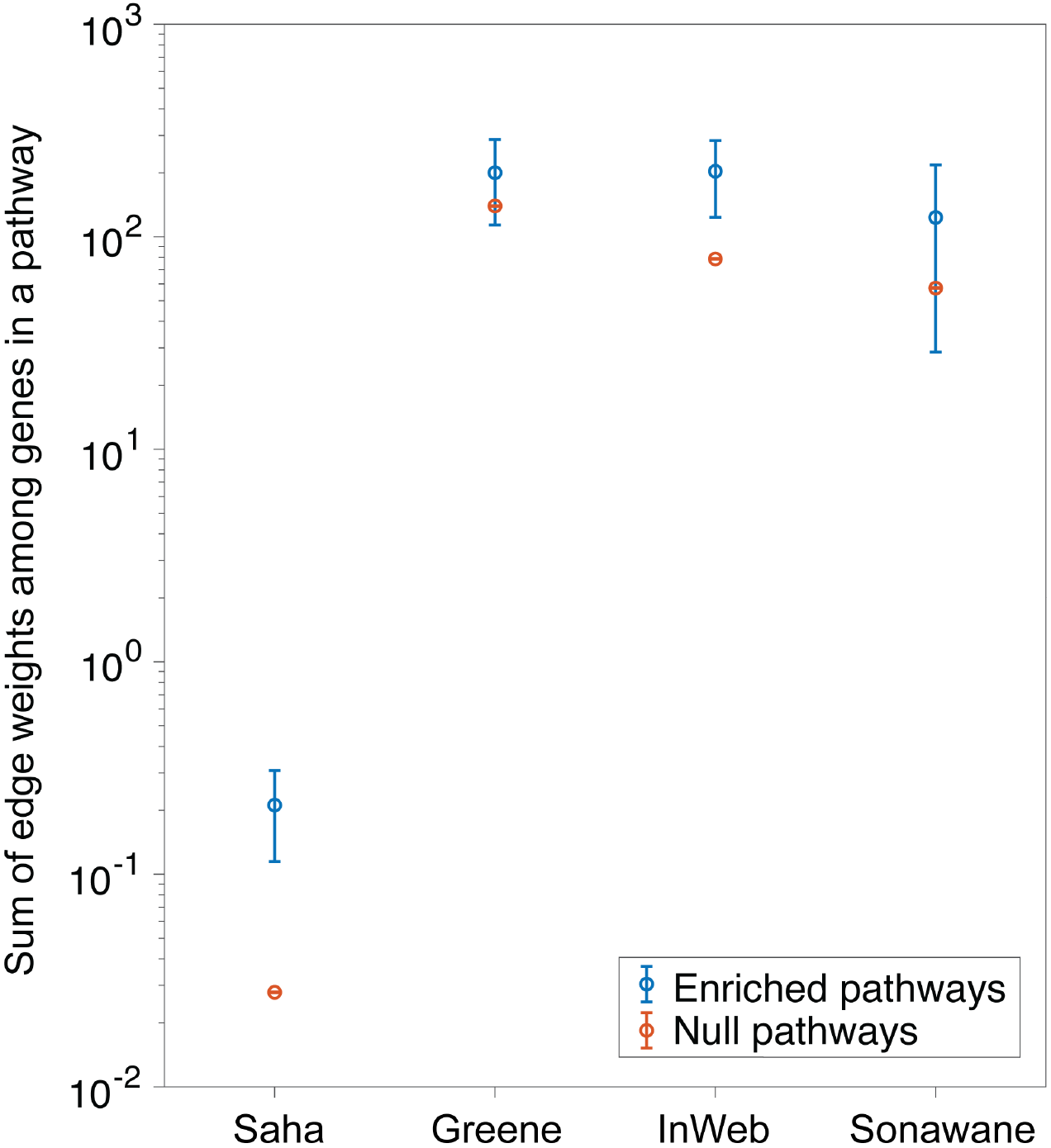
Genes in enriched pathways have high network connectivity. For each of four networks, we report the sum of edge weights in the network between genes in the pathway, averaged across 141 enriched pathways. For comparison purposes, we report the same quantity averaged across 10,000 null pathways with the same number of genes. Error bars represent 95% confidence intervals (smaller than data points for null pathways). Numerical results, and analogous results for network connectivity between a pathway and interacting genes outside the pathway, are reported in Table S22.

We repeated the S-LDSC analysis (Figure 3b-c) using new pathway+network annotations constructed using a random-forest classifier, Quack (Li et al., 2018), that identifies new candidate genes that have similar topological patterns and network centrality metrics as genes in the input pathway. Because genes in enriched input pathways have high network connectivity (Figure 4), this is closely related to our primary strategy of defining pathway+network annotations based on genes with high network connectivity to genes in enriched input pathways. Indeed, Quack annotations were highly correlated with our main pathway+network annotations (*r* =0.50-0.67; Table S12), and produced S-LDSC results similar to our main analysis (Figure 3b-c), including significant enrichments for all four networks but non-significant *τ*\* conditional on the baseline-LD model (Table S23).

## Discussion

We analyzed 42 diseases and complex traits (average *N*=323K) to show that genes with high network connectivity are enriched for disease heritability, but that it is critical for gene network and pathway analyses to account for known functional annotations, such as those from our baseline-LD model (Gazal et al., 2017). First, in analyses of pathway annotations, we identified 156 pathway-trait pairs with significant heritability enrichment after conditioning on the baseline-LD model, a stringent step that caused a majority of pathway-trait pairs reported in recent studies to become non-significant in our analyses. Second, we determined that network annotations based on closeness centrality, a measure of network connectivity, are strongly enriched for disease heritability, but that these enrichments were fully explained by annotations from the baseline-LD model. Third, for each of the 156 significant pathway-trait pairs, we determined that pathway+network annotations constructed from genes with network connectivity to the input pathway were strongly enriched for the corresponding traits, but that once again these enrichments were largely explained by annotations from the baseline-LD model.

Our findings have important ramifications for studies connecting gene networks and pathways to disease (Köhler et al., 2008; Vanunu et al., 2010; Segrè et al., 2010; Lee et al., 2011; Califano et al., 2012; Pers et al., 2015; de Leeuw et al., 2015; Taşan et al., 2015; Greene et al., 2015; de Leeuw et al., 2016; Marbach et al., 2016; Boyle et al., 2017; Peters et al., 2017; Cowen et al., 2017; Pardiñas et al., 2018; Yoon et al., 2018; Wray et al., 2018; Savage et al., 2018; Zhu and Stephens, 2018; Nagel et al., 2018). Specifically, it is important to account for known functional annotations when seeking to elucidate biological mechanisms. For some methods, such as S-LDSC (Finucane et al., 2015; Boyle et al., 2017; Pardiñas et al., 2018), it is straightforward to incorporate known functional annotations such as those from the baseline-LD model (Gazal et al., 2017), and we emphasize the importance of doing so. For other methods, it is of high interest to investigate how functional annotations could be incorporated. More generally, it is of broad interest to re-assess previously reported results while accounting for known functional annotations; for example, this could be achieved by running S-LDSC both with and without incorporating functional annotations from the baseline-LD model.

We note several limitations of our work. First, S-LDSC is not well-suited to analysis of annotations spanning a very small proportion of the genome (Finucane et al., 2015) and does not model sparsity in trait effect sizes, potentially explaining why we did not identify enriched pathways for 8 traits that are less polygenic (O’Connor et al., 2018) (e.g. age at menopause, balding, hair color, sunburn). Nonetheless, our main results attained high statistical significance. Second, we did not explicitly compare S-LDSC to other methods. However, previous work suggests that S-LDSC compares favorably to other gene set enrichment methods, both in simulations and in analyses of real traits (Finucane et al., 2018). Third, interpretation of pathway-trait enrichments is complicated by the possibility that enrichment signals may be driven by a small number of highly significant genes (Zhu and Stephens, 2018). Indeed, when we repeated our analyses excluding from each enriched pathway all genes harboring genome-wide significant associations from the corresponding trait (average of 2 genes removed per pathway-trait pair; see Methods), only 53 pathway-trait pairs remained significant (Table S24). However, we verified that repeating our main pathway+network analyses using these 53 pathway-trait pairs (excluding genes implicated by GWAS) produced similar conclusions (Table S25). Fourth, gene networks may include false-positive interactions, even after correcting for technical confounding (Parsana et al., 2017; Saha et al., 2017; Boyle et al., 2018). However, our results are consistent across four published gene networks (Greene et al., 2015; Saha et al., 2017; Li et al., 2017; Sonawane et al., 2017), suggesting that false-positive interactions are unlikely to substantially impact our results. Fifth, inferences about components of heritability can potentially be biased by failure to account for LD-dependent architectures (Speed et al., 2012; Yang et al., 2015; Gazal et al., 2017; Speed et al., 2017). All of our main analyses used the baseline-LD mode, which includes 6 LD-related annotations (Gazal et al., 2017). The baseline-LD model is supported by formal model comparisons using likelihood and polygenic prediction methods, as well as analyses using a combined model incorporating alternative approaches (Gazal et al., 2018); however, there can be no guarantee that the baseline-LD model perfectly captures LD-dependent architectures. Finally, although we identified many significantly enriched pathways, our results for network and network+pathway annotations represent a negative result for efforts to improve upon the baseline-LD model (Gazal et al., 2017). On the other hand, our results emphasize the importance of accounting for known functional annotations in network and pathway analyses.

## Methods

### Pathway annotations

We considered 18,119 gene sets (pathways) from five sources: 2,118 biological pathways from the BioSystem (BS) database (Geer et al., 2010) (which includes pathways from BioCyc (Caspi et al., 2018), Kyoto Encyclopedia of Genes and Genomes (KEGG) (Kanehisa et al., 2017), Pathway Interaction Database (PID) (Schaefer et al., 2009), REACTOME (Fabregat et al., 2018), WikiPathways (Pico et al., 2008)), 1,927 biological pathways from the Pathway Commons (PC) database (Cerami et al., 2011) (which includes pathways from HumanCyc (Romero et al., 2004), Integrating Network Objects with Hierarchies (INOH) (Yamamoto et al., 2011), KEGG (Kanehisa et al., 2017), PANTHER (Mi and Thomas, 2009), PID (Schaefer et al., 2009), REACTOME (Fabregat et al., 2018), SMPDB (Jewison et al., 2014), NetPath (Kandasamy et al., 2010)), 7,209 protein-protein interaction gene sets from the InWeb database (Li et al., 2017), 3,903 mouse phenotype gene sets from the Mouse Genome Informatics (MGI) database (Eppig et al., 2014)(i.e. sets of genes whose orthologs are associated to mouse phenotypes), and 2,961 gene ontology gene sets from the Genome Ontology (GO) database (Ashburner et al., 2000). This set of pathways, which contain at least 10 genes and at most 500 genes (consistent with previous studies (Pers et al., 2015; Zhu and Stephens, 2018), significantly overlap with mSigDB (Liberzon et al., 2011) and pathways analyzed in a previous study (Pers et al., 2015).

For each of 18,119 pathways, we constructed a pathway annotation by annotating a value of 1 for variants around the protein-coding genes in a given pathway (+/− 100kb) and 0 for all other variants. We applied S-LDSC on 18,119 pathway annotations across 42 independent diseases and complex traits, conditional on the baseline-LD model (v1.1) (Gazal et al., 2017) and all-genes annotations (a value of 1 for variants around 23,987 protein-coding genes with a 100kb window). We removed 129 pathway-trait pairs whose annotated SNPs are less than 0.02% of the reference genome (European samples from the 1000 Genomes Project; see URLs) as S-LDSC is not well-equipped for annotations that span very small proportion of the genome.

For each of 760,869 pathway-trait pairs, we assessed the statistical significance of the pathway annotations based on global FDR < 5% on the pathway annotation’s standardized effect size (*τ*\*) p-value. We note controlling FDR for each trait and for all traits did not make a major difference in the number of identified enriched pathway-trait pairs. We further note that our choice of two-tailed test on the significance is conservative, partially attributing to the reduced number of significantly associated pathway-trait pairs. Among significantly enriched pathway-trait pairs. we calculated a pairwise correlation for every pair of annotations and retained the more significant pathway for correlated pathways with r ≥ 0.5 (as in a previous study (Li et al., 2018)). If correlated pathways were enriched for different traits, we retained both pathways.

For each of 156 significantly enriched pathway-trait pairs, we also constructed pathway annotations excluding genes implicated by GWAS. First, we downloaded all GWAS associations from the GWAS Catalog (see URLs); we restricted to significant associations (p-value ≤ 5e-8). For each of the 34 traits, we defined genes implicated by GWAS by including genes mapped to the lead SNP; if the SNP was intergenic, we included the nearest upstream and downstream genes. (We note that the nearest gene might not be the correct target gene (Gusev et al., 2016).) Then, for each of the 156 enriched pathway-trait pairs, we removed trait-specific GWAS significant genes (5% of the genes, on average across 156 pathway-trait pairs) and rebuilt our pathway annotations with a 100kb window. For pathways significant for multiple traits, we built unique pathway annotations excluding genes implicated by GWAS for the corresponding traits. For each of 156 pathway-trait pairs, genes after excluding GWAS significant genes are provided in Table S4.

### Network annotations

We downloaded each of four gene networks (Greene et al., 2015; Saha et al., 2017; Li et al., 2017; Sonawane et al., 2017) and formatted as a two-dimensional matrix with three columns: gene1, gene2, and interaction score (see URLs). We used ENTREZ ID for the gene identifier, so converted other gene identifiers provided by the authors as needed. For each network, we constructed seven different probabilistic annotations based on the following network connectivities (centralities), which we computed using Graph-tool (see URLs):

Let G be a weighted, undirected graph with a set of vertices (V) and a set of edges (E).

*Closeness centrality (v):* how close gene v is to all other genes in the network. It is defined as 1/∑_*v*,*v*≠*w*_ *d*_*vw*_ where *d*_*vw*_ is the weighted distance from v to w. If there is no path between two genes, the distance of zero is used.

*Degree (v):* the number of vertices connected to v; that is the number of neighboring genes for each gene in the graph.

*Maximum edge weight (v):* the maximum of the weights of all edges connected to v.

*Sum edge weight (v):* the sum of the weights of all edges connected to v.

*Betweenneses centrality (v):* The number of shortest paths that pass through gene v. It is defined as ∑*σ*_*uw*_(*v*)/*σ*_*uw*_ where *σ*_*uw*_ (*v*) is the number of shortest paths from node u to w that pass through v, and *σ*_*uw*_ is the total number of shortest paths from u to w.

*Eigenvector centrality (v):* Intuitively, the eigencentrality of v is proportional to the sum of the centralities of its neighbors (Özgür et al., 2008). It is defined as the solution of of *Ax* = λ*x*, where *x* is the eigenvector of the weighted adjacency matrix *A* with the largest eigenvalue *λ*.

*Pagerank (v):* Pagerank is similar to eigenvector centrality, except it contains a damping factor, the probability that the person who randomly visit genes will continue, under the assumption that more important genes are more likely to receive more visits. It is defined as (1 – *d*)/*N* + *d*∑_*u*∈*N*(v)_ *Pagerank*(*u*)/*d*^+^(*u*) where *d* is a damping factor, *N*(*v*) are the in-neighbors of v, *d*^+^(*u*) is the out-degree of u. Because G is an undirected graph, pagerank treats it as a directed graph by making edges bidirectional.

After computing network connectivities for all protein-coding genes that exist in a network, we linearly transformed scores to lie between 0 to 1 and annotated variants around genes (+/− 100kb). When a variant is spanned by multiple genes with the 100kb window, we assigned the maximum connectivity score.

### Pathway + Network annotations

We constructed pathway+network annotations, specific to an input pathway and a gene network. For each of 141 enriched pathways, we first constructed an adjacency matrix by mapping genes in a pathway (“core genes”) to a gene network and identifying the set of neighboring genes (“peripheral genes”), using Graph-tool (see URLs). In the adjacency matrix, a vector represents an ENTREZ ID of a gene in a pathway, an ENTREZ ID of a neighboring gene, and the interaction score (e.g. posterior probability of these two genes). Using this adjacency matrix representing a subnetwork of core and perihpheral genes that appear in the network, we score both core and peripheral genes based on closeness centrality. We use the inverse of the interaction score as the cost when computing closeness centrality. Using the set of core and peripheral genes along with the linearly transformed connectivity scores that lie from 0 to 1, we annotated variants around genes (+/− 100kb). When a variant is spanned by multiple genes with the 100kb window, we assigned the maximum connectivity score.

### Excess overlap between annotations

We calculated excess fold overlap between two annotations (annot 1 and annot 2) as the fraction of overlap between the annotations divided by the expected amount of overlap by chance, computed as the following: excess overlap(annot1, annot2) =

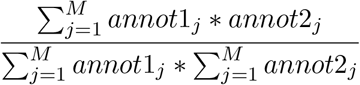

where M is the total number of SNPs (5,961,159). Annotations with no excess overlap are assigned a value of 1; when there is excess overlap, the excess fold overlap is > 1.

### Network connectivity of a pathway

Given a pathway and a network, we quantified how tightly connected the pathway is in the network, compared to a null pathway. Let P be a set of genes in the pathway that exist in the network. Let Q be a set of neighboring genes; that is, genes in P are connected with genes in Q with at least one edge, in the network. We calculated three metrics: (1) size of Q (number of neighboring genes), (2) sum of edges among genes in P, and (3) sum of edges between genes in P and genes in Q. We did not consider genes in the pathway that are not coding or do not appear in the network.

For each of 141 enriched pathways, we constructed a corresponding null pathway as follows: (1) randomly choose a pathway from the full set of 18,119 pathways (Table S1), (2) randomly choose a gene from the sampled pathway, and (3) repeat (1) and (2) N times, where N is the number of genes in the pathway. For each of four gene networks analyzed, we repeated this procedure 10,000 times and reported three connectivity metrics of enriched and null pathways (for null, the mean of 10,000 permutations is reported).

### Set of 42 independent traits

Analogous to a previous study (Hormozdiari et al., 2018), we considered 89 GWAS summary association statistics, including 34 traits from publicly available sources and 55 traits from the UK Biobank (up to N = 459K); summary association statistics were computed using BOLT-LMM v2.3 (Loh et al., 2015, 2018). Among 47 summary statistics with z-scores of total SNP heritability of at least 6 (computed using S-LDSC with the baseline-LD model), we further removed 5 summary statistics that have genetic correlation of at least 0.9 (computed using cross-trait LDSC (Bulik-Sullivan et al., 2015)). Whenever applicable, meta-analysis across 42 independent traits (Table S2), whose GWAS summarystatistics are publicly available (see URLs), was performed using a random-effect meta-analysis using the R package rmeta.

### Set of 10 blood-related and 8 brain-related traits

We analyzed ten independent blood-and autoimmune-related traits: Crohn’s disease (Jostins et al., 2012), rheumatoid arthritis (Okada et al., 2014), and ulcerative colitis (Jostins et al., 2012) from publicly available datasets and eczema, autoimmune diseases. eosinophil count, platelet count, red blood cell count, white blood cell count, and red blood cell distribution width from the UK Biobank.

We analyzed eight independent brain-related traits: autism spectrum (of the Psychiatric Genomics Consortium et al., 2013), depressive symptoms (Okbay et al., 2016), and schizophrenia (Ripke et al., 2014) from publicly available datasets, and age at menarche, body mass index, neuroticism, smoking status, and years of education from the UK Biobank. We selected these traits from 42 independent traits we analyzed, as inferred from heritability enrichment of tissue-specific gene expression and chromatin annotations (Finucane et al., 2015, 2018) and eQTL annotations (Hormozdiari et al., 2018). We additionally considered autism spectrum based on previous studies (Fatemi et al., 2009; Cellot and Cherubini, 2014) that the brains of subjects with autism have altered expression.

## Data access

All pathway, network, and pathway+network annotations and LD scores are publicly available (see URLs).

## URLs

Pathway, network, and pathway+network annotations: https://data.broadinstitute.org/alkesgroup/LDSCORE/Kim_pathwaynetwork

S-LDSC software: https://github.com/bulik/ldsc

baselineLD annotations: https://data.broadinstitute.org/alkesgroup/LDSCORE/

Saha transcriptome-wide networks: http://gtexportal.org

Greene tissue-specific co-expression networks: http://hb.flatironinstitute.org/

InWeb protein-protein interaction network: https://www.intomics.com/inbio/map

Sonawane gene regulatory networks: https://zenodo.org/record/838734

GTEx (Release v6, dbGaP Accession phs000424.v6.p1): http://www.gtexportal.org.

GWAS Catalog (Release v1.0): https://www.ebi.ac.uk/gwas.

1000 Genomes Project Phase 3 data: ftp://ftp.1000genomes.ebi.ac.uk/vol1/ftp/release/20130502

PLINK software: https://www.cog-genomics.org/plink2

BOLT-LMM software: https://data.broadinstitute.org/alkesgroup/BOLT-LMM

BOLT-LMM summary statistics for UK Biobank traits: https://data.broadinstitute.org/alkesgroup/UKBB

UK Biobank: http://www.ukbiobank.ac.uk/

UK Biobank Genotyping and QC Documentation: http://www.ukbiobank.ac.uk/wp-content/uploads/2014/04/UKBiobank_genotyping_QC_documentation-web.pdf

Graph-tool: https://graph-tool.skewed.de

## Acknowledgements

We are grateful to Manolis Kellis, Yakir Reshef, Evan Boyle, April Kim, and Marieke Kuijjer for helpful discussions. This research was funded by NIH grants U01 HG009379, R01 MH109978, R01 MH101244 and R01 MH107649. This research was conducted using the UK Biobank Resource under Application 16549.

## Contributions

S.S.K. and A.L.P. conceived the project. S.S.K. performed the experiment. S.S.K. and C.D. analyzed data. A.L.P. supervised analyses. S.S.K. and A.L.P. wrote the manuscript with assistance from all authors.

## Supplementary figures

**Figure S1.**
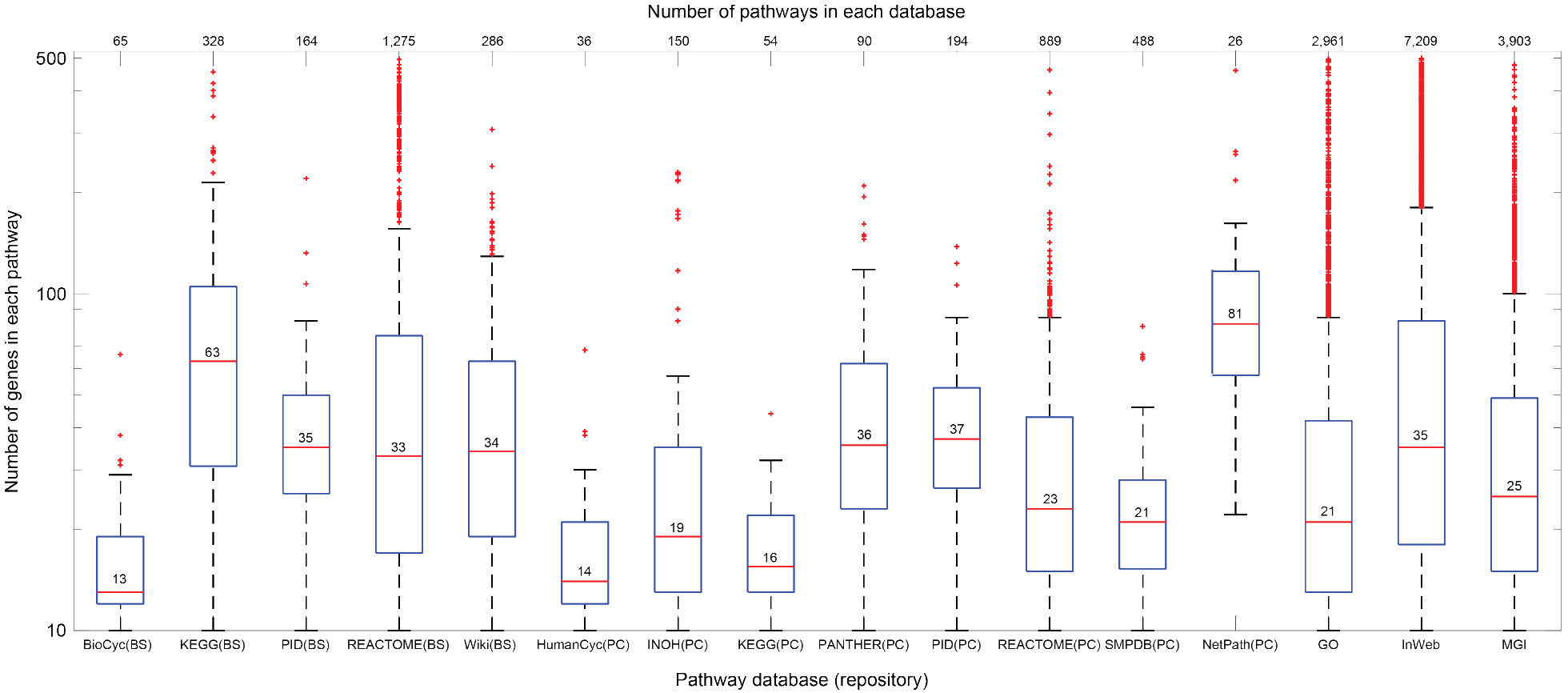
Summary of pathways analyzed. We analyzed 18,119 pathways, each consisting of at least 10 and at most 500 coding genes. Number in the boxplot represents the median number of genes for each database. Number of pathways in each database is shown on the top of the figure. BS: NCBI BioSystems. PC: PathwayCommons. MGI: Mouse Genome Informatics. GO: Genome Ontology. InWeb: InWeb protein-protein interactions. See Table S1 for a description of 18,119 pathways analyzed.

**Figure S2.**
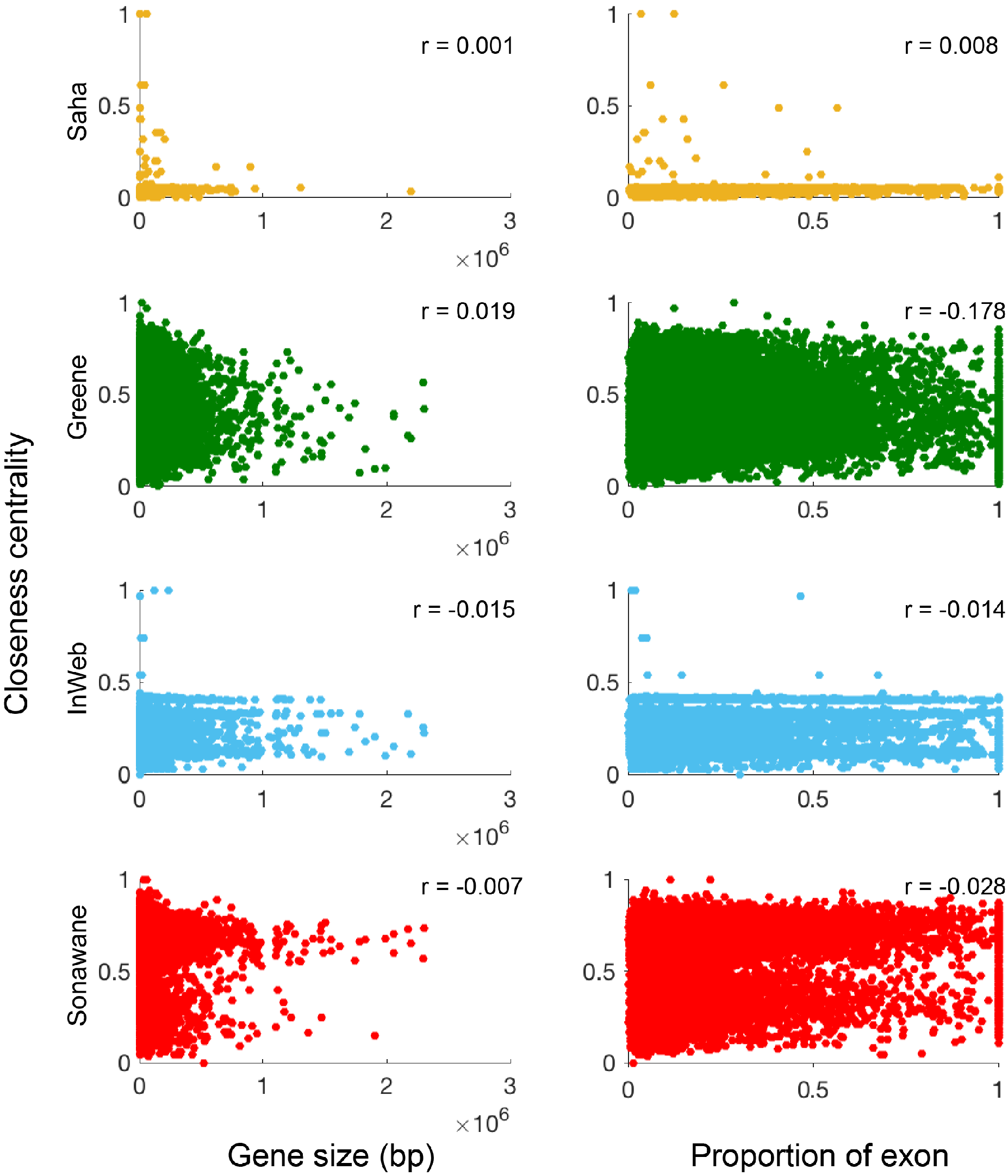
Closeness centrality is independent from the gene size and the proportion of exon. For each of four network annotations, we computed a Pearson correlation between probabilistic annotation values and (1) gene size and (2) proportion of exon. We calculated the proportion of exon as the size of coding regions (bp) that lie inside the gene divided by the size of the gene, defined as (transcription stop-transcription start).

## Supplementary tables

See Excel file for all supplementary tables. Titles and captions are provided below.

**Table S1. List of 18,119 pathways analyzed.** For each pathway, we report a pathway ID, pathway description, database, ENTREZ IDs for genes, and the number of protein-coding genes.

**Table S2. List of 42 independent diseases and traits analyzed.** For each trait, we report a trait identifier, trait description, reference, sample size, and heritability z-score. We selected these 42 traits based on a heritability z-score > 6 (see Methods). We further indicated brain or blood-related traits.

**Table S3. S-LDSC results of all pathway-trait pairs** We applied S-LDSC to 760,869 pathway-trait pairs, conditioning on all-genes annotation and the baseline-LD model. For each pathway-trait pair, we report a proportion of SNPs, enrichment, and *τ*.

**Table S4. S-LDSC results for 156 significantly enriched pathway-trait pairs.** For each significantly enriched pathway-trait pair, we report a proportion of SNPs, enrichment, and a *τ*∗ The 8 significant pathway-trait pairs were reported in previous genetic studies: “pathways in cancer” for height (Wood et al., 2014); “neuropeptide hormone activity” for BMI (Locke et al., 2015); “immune response” for both Crohn’s disease and ulcerative colitis (Jostins et al., 2012); “T-cell receptor,” “abnormal T-cell physiology,” and “cytokine-mediated signaling pathway” for rheumatoid arthritis (Okada et al., 2014); “absent corpus callosum” for years of education. (Okbay et al., 2016)

**Table S5. Average gene size of annotations.** For all-genes annotation, all pathways, pathway, network, and pathway+network, and Quack annotations, we report an average size of genes (and its standard deviation) and an average number of genes.

**Table S6. Heritability enrichment of enriched pathway-trait pairs.** We meta-analyzed (a) 156 enriched pathway-trait pairs; (b) 13 enriched pathway-trait pairs for ExAC, Cassa, and Samocha gene sets; (c) 169 enriched pathway-trait pairs (a and b combined). In each case, we report meta-analyzed enrichments and *τ*∗.

**Table S7. S-LDSC results of 195 pathway-trait pairs from previous pathway enrichment studies.** We applied S-LDSC to 95 pathway-trait pairs from five previous genetic studies (Boyle et al., 2017; Pardiñas et al., 2018; Wray et al., 2018; Savage et al., 2018; Nagel et al., 2018) and 100 from an unpublished study (Zhu and Stephens, 2018) (a) conditioning on the baseline-LD model and all-genes annotation and (b) conditioning on all-genes annotation only. In each pathway-trait pairs for each case, we report a proportion of SNPs, enrichment, and *τ*.

**Table S8. S-LDSC results of 13 enriched pathway-trait pairs for ExAC, Cassa, Samocha gene sets.** For each of 13 enriched pathway-trait pairs, we report a proportion of SNPs, enrichment, and *τ*.

**Table S9. Summary of gene networks analyzed.** We report the number of genes and the number of edges for each of four networks (InWeb, Saha, Sonawane, Greene).

**Table S10. Correlation of network annotations with baseline-LD model annotations.** We report the Pearson correlation between baseline-LD model annotations and (a) Saha network annotations of different centralities, (b) Saha network annotations of different tissues, (c) Greene network annotations of different centralities, (d) Greene network annotations of different tissue, (e) InWeb network annotations of different centralities, (f) Sonawane network annotations of different centralities, and (g) Sonawane network annotations of different tissue.

**Table S11. Excess fold overlap of pathway and network based annotations with baseline-LD model annotations.** We report the excess fold overlap between baseline-LD model annotations and pathway, network, pathway+network, Quack, and all-genes annotations.

**Table S12. Correlation of pathway and network based annotations with baseline-LD model annotations.** We report the Pearson correlation between baseline-LD model annotations and pathway, network, pathway+network, Quack, and all-genes annotations.

**Table S13. Excess fold overlap / correlation among functional annotations from the baseline-LD model.** We report (a) excess fold overlap and (b) correlation among functional annotations from the baseline-LD model.

**Table S14. Heritability enrichment of network annotations.** For each of 4 network annotations, we report meta-analyzed enrichments and *τ*∗ across 42 independent traits.

**Table S15. Heritability enrichment of network annotations conditioning on one annotation from the baseline-LD at a time.** For (a) Saha, (b) Greene, (c) InWeb, (d) Sonawane networks, we meta-analyzed network annotations across 42 independent traits, conditioning on one annotation from the baseline-LD model at a time. We report meta-analyzed enrichments and *τ*∗ across 42 independent traits.

**Table S16. Heritability enrichment of network annotations using relevant tissues for brain-related and blood-related traits.** For (a) 8 brain-related traits and (b) 10 blood-related traits, we report meta-analyzed enrichments and *τ*∗.

**Table S17. Heritability enrichment of network annotations using 6 other network centrality metrics.** For (a) Saha, (b) Greene, (c) InWeb, (d) Sonawane networks, we constructed network annotations based on 6 other network centrality metrics (see Methods) and meta-analyzed results across 42 independent traits. We report meta-analyzed enrichments and *τ*∗.

**Table S18. Heritability enrichment of network annotations with different window sizes.** Instead of 100kb windows around genes, we added (a) 10kb or (b) 1Mb windows around genes when constructing network annotations. We report meta-analyzed enrichments and *τ*\* across 42 independent traits.

**Table S19. Heritability enrichment of DSD-network annotations.** We applied the diffusion state distance (DSD) algorithm (Cao et al., 2013) to transform gene networks’ edge weights with a random walk (k = 5). Then, we constructed network annotations by re-computing closeness on DSD-transformed networks and meta-analyzed results across 42 independent traits. We report meta-analyzed enrichments and *τ*∗.

**Table S20. Heritability enrichment of pathway+network annotations.** We meta-analyzed 156 pathway-trait pairs (122 for Saha, which has less pairs as all genes in some pathways do not exist in the Saha network). We report meta-analyzed enrichments and *τ*∗ across 156 pathway-trait pairs (122 for Saha).

**Table S21. Heritability enrichment of pathway+network annotations conditioning on one annotation from the baseline-LD at a time.** For (a) Saha, (b) Greene, (c) InWeb, (d) Sonawane networks, we constructed an average annotation across 156 pathway+network annotations and meta-analyzed averaged pathway+network annotations across 42 independent traits, conditioning on one annotation from the baseline-LD model at a time. We report meta-analyzed enrichments and *τ*∗ across 42 independent traits.

**Table S22. Network connectivity of enriched pathways and null pathways.** Using (a) sum of edge weights or (b) number of edges as network connectivity metrics, we report the number of interacting genes and network connectivity between genes in a pathway (core edges) and genes outside the pathway (peripheral edges). We constructed null pathways in two ways: (1) each gene sampled from a randomly chosen pathway or (2) each gene randomly sampled from all protein-coding genes.

**Table S23. Heritability enrichment of Quack annotations.** We applied the Quack random-forest classifier algorithm (Li et al., 2018). We used 18,119 pathways as a training data and applied Quack to four gene networks. We used the output of Quack to construct Quack pathway+network annotations (with 100kb window), applied S-LDSC, and meta-analyzed across 156 pathway-trait pairs (122 for Saha). We report meta-analyzed enrichments and *τ*∗.

**Table S24. S-LDSC results of 156 enriched pathway-trait pairs excluding genes implicated by GWAS.** We removed genes implicated by previous GWAS studies (see Methods; average of 5% of genes (2 genes) removed) and applied S-LDSC conditional on all-genes and baseline-LD model annotations. For each pathway-trait pair, we report a proportion of SNPs, enrichment, *τ*∗, and the number of genes in a pathway excluding GWAS significant genes.

**Table S25. Heritability enrichment of 53 pathway+network annotation using pathways excluding genes implicatd by GWAS.** From 53 significant pathway-trait pairs after excluding GWAS significant genes, we constructed pathway+network annotations and meta-analyzed results across 53 pathway-trait pairs (40 for Saha). We report meta-analyzed enrichments and *τ*∗.

